# H&Enium, Applying Foundation Models to Computational Pathology and Spatial Transcriptomics to Learn an Aligned Latent Space

**DOI:** 10.1101/2025.07.22.665986

**Authors:** Marc Glettig, Tim Ehrensperger, Josephine Yates, Valentina Boeva

**Affiliations:** Department of Computer Science, Eidgenössische Technische Hochschule Zürich, ETHZ, Zü rich, Switzerland; Dana Farber Cancer Institute, DFCI, Boston, United States of America

## Abstract

Bridging the gap from transcriptomic to imaging data at single-cell resolution is essential for understanding tumor biology and improving cancer diagnostics. Spatial transcriptomics enables mapping gene expression onto H&E images of segmented single cells, but remains limited by cost and throughput. We introduce H&Enium, a contrastive alignment framework that projects image and gene expression embeddings from foundation models into an aligned latent space using projection heads and a novel soft alignment target. This alignment enriches image-derived embeddings with transcriptomic context improving downstream tasks such as cell type classification and gene expression prediction. Additional evaluations on independent pathology datasets demonstrate superior generalization of our aligned representations over unaligned baselines. Our method offers a scalable path to enhance the utility of standard H&E imaging in both research and clinical settings.

## 1. Introduction

Cancer remains a leading cause of death worldwide, with its complexity posing significant challenges to effective treatment. Consequently, advancing research to improve diagnosis and therapeutic strategies is crucial. Recently, large pretrained machine learning models have shown great promise in cancer genomics and pathology, with clinically approved applications emerging in computational pathology (Campanella et al., 2019; Yates & Allen, 2025).

Foundation models, leveraging extensive datasets to capture complex patterns, have become particularly influential in digital pathology. They have successfully addressed tasks such as tissue classification, biomarker detection, and gene expression prediction. Example models include UNI (Chen et al., 2024) and CONCH (Lu et al., 2024). Additionally, CONCH uses a bi-modal training procedure, including pathology reports, to improve image embeddings. Similarly, transcriptomics foundation models have enabled significant progress in the characterization of cellular heterogeneity and gene expression dynamics. Notable models include CellPLM (Wen et al., 2023) and scGPT (Cui et al., 2024) which are pretrained on single-cell RNA sequencing data using masked gene prediction objectives, analogous to masked language modeling in NLP.

With the advent of spatial transcriptomics, recent studies have aimed to align imaging and transcriptomic modalities using contrastive training frameworks inspired by CLIP (Radford et al., 2021). Methods such as BLEEP (Xie et al., 2023), ST-Align (Lin et al., 2024), and PathOmCLIP (Lee et al., 2024) improved spatial modality alignment, primarily at the spatial-spot resolution. BLEEP uses soft targets to account for input embedding similarities. ST-Align introduced patch-level foundation models to embed the two input modalities. Finally, PathOmCLIP (Lee et al., 2024) adds the usage of a local transformer to allow for the incorporation of neighboring patch embeddings, thus improving spatial context and alignment. While recent advancements have improved the alignment between H&E imaging and spatial transcriptomics at the spot level, they fall short at singlecell resolution, the scale at which key biological insights emerge from interactions between individual cells and their surrounding microenvironments.

In this work, we introduce **H&Enium**, a self-supervised model leveraging pathology and transcriptomics foundation models to learn aligned latent representations of single cells across both modalities. Using Xenium (Janesick et al., 2023) spatial transcriptomics slides, we demonstrate that readily available H&E foundation model embeddings, although trained only on patch level data, substantially improve cell type classification accuracy from H&E images. This performance can be further improved by the aligned latent space. Aligned cell embeddings derived from imaging outperform zero-shot foundation model embeddings, enhancing cell typing accuracy by more than 16% and gene expression prediction by more than 10% consistently across samples. The ability to accurately predict cell types directly from H&E whole-slide images unlocks the potential for spatial analyses in existing large-scale pathology datasets. This will significantly advance our understanding of the tumor microenvironment and tumor biology.

## 2. Methods

### 2.1. H&Enium Architecture

Each (single) cell is represented by the tuple

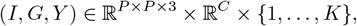

where *I* is the *P × P* H&E-stained image patch, *G* the *C*-dimensional gene-expression vector, and *Y* the cell-type label out of *K* classes.

Figure 1 illustrates the H&Enium architecture. Given an image patch *I* and a gene vector *G* of a single cell, we first extract frozen embeddings via a pathology foundation model and a transcriptomic foundation model, respectively:

**Figure 1.**
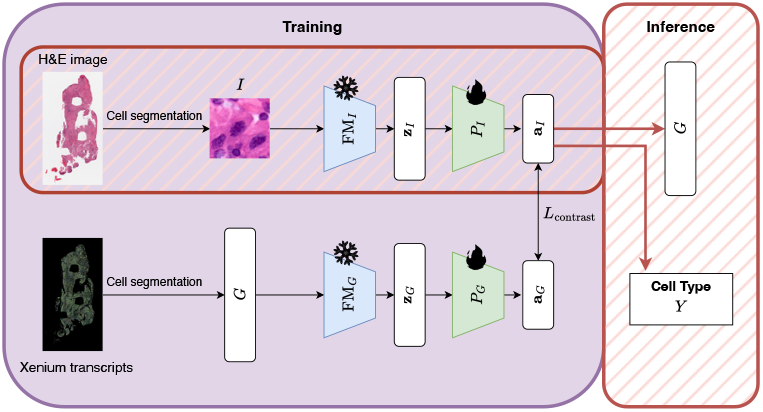
Overview of the H&Enium single-cell alignment model architecture. During training, all foundation models (FMs) remain frozen, while projection heads *P*_*I*_ and *P*_*G*_ are jointly trained using the contrastive loss *L*_contrast_.

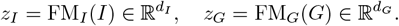

Where *d*_*I*_ and *d*_*G*_ are the foundation model embedding dimensions. For a batch of *B* cells we write 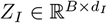 and 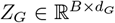. Two projection heads *P*_*I*_, *P*_*G*_ are then jointly trained to align these embeddings:

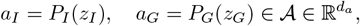

or in batch form 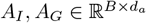.

We align image and gene embeddings via a contrastive loss that pulls matching pairs together and pushes non-matching pairs apart. To compute the loss, we first perform row-wise L2-normalization in the aligned space and then calculate the cosine similarity matrix between gene and image embeddings as follows: *S* = *cossim*(*A*_*G*_, *A*_*I*_) ∈ ℝ^*B×B*^, where each entry *S*_*ij*_ = ⟨ *a*_*G,i*_, *a*_*I,j*_ ⟩ measures the similarity between gene embedding *i* and image embedding *j*.

Let *T* ∈ [0, 1]^*B×B*^ be the target similarity matrix, where each entry *T*_*ij*_ encodes the desired pairing strength between cell *i* and cell *j*. In our contrastive loss, we directly compare each predicted similarity *S*_*ij*_ to its target *T*_*ij*_ via a soft crossentropy, formally defined in Appendix A, and apply it in both directions: *L*_gene_ = SoftCE(*S, T*), and *L*_image_ = SoftCE(*S*^⊤^, *T* ^⊤^).

The final contrastive loss

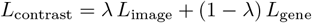

uses *λ* ∈ [0, 1] to balance the contributions of each alignment direction.

We evaluate three different targets *T*, named CLIP, BLEEP and BLEEP_input_. CLIP is a (one-hot) diagonal target used in Radford et al. (2021), BLEEP is a soft target derived from the aligned space 𝒜 defined by Xie et al. (2023). BLEEP_input_ is our newly introduced soft target based on the pre-projection embeddings *z*. Details on target definitions can be found in Appendix B. As projection heads, we use a simple MLP. For details on the projection heads and training procedure, refer to Appendix C.

### 2.2. Downstream Task Modeling

We assess embedding quality on cell type classification and gene expression prediction using the frozen foundation model embeddings *z*_*I*_ and *z*_*G*_ and their aligned counterparts *a*_*I*_ and *a*_*G*_.

#### 2.2.1. Cell type classification

We train L2-penalized logistic regressions with balanced class weights and the L-BFGS solver (up to 5,000 iterations) on both frozen foundation model embeddings (*z*_*I*_, *z*_*G*_) and aligned embeddings (*a*_*I*_, *a*_*G*_). We report accuracy, balanced accuracy (BAC), and F1 score (F1).

#### 2.2.2. Gene expression prediction

We predict gene expression solely from the image-based embeddings *z*_*I*_ and *a*_*I*_. Following (Jaume et al., 2024), we filter out genes expressed in less than 10% of cells, normalize counts to counts-per-million (CPM), apply log 1*p* transformation, and select the top 50 most variable genes. To predict the log-CPM values of the 50 most variable genes, we standard-scale each embedding vector, apply PCA (keeping 64 principal components if the embedding dimension is larger than 64), and then fit a Ridge regression model.

We report Pearson correlation (PCC) and Relative Variance Distance (RVD), explained in Appendix D.

### 2.3. Baselines

As a naive baseline, we employ a majority label classifier that consistently predicts the most common label in the training set. For a more informed comparison, we construct a morphological baseline using geometric features—such as cell and nuclear area, perimeter, and shape descriptors—extracted from Xenium’s cell segmentations. This allows us to assess the added value of representations from pathology foundation models over handcrafted morphological cues (see Appendix E).

## 3. Results

### 3.1. Dataset

To train and test H&Enium, we used publicly available tumor spatial transcriptomic slides generated on the 10x Genomics Xenium platform. We worked with three distinct slides from pancreas and breast, capturing over one million individual cells. We obtained cell type labels *Y* via expert annotation based solely on Xenium single-cell gene expression data. The labels *Y* correspond to the four PanNuke (Gamper et al., 2020) classes Connective, Inflammatory, Neoplastic, and Epithelial.

Additionally, we evaluated our approach on the much smaller out-of-sample PanNuke H&E test set (Gamper et al., 2020). No corresponding gene expression data is available for the PanNuke dataset, i.e. pathologists annotated the cell type labels by eye. For additional information on the datasets or visual context, refer to Appendix F.

### 3.2. Preprocessing

For each nucleus centroid, we extract one square image patch *I* of *P* = 224 pixels centered on the centroid, which matches the input size for UNI2 and CONCH, the pathology foundation models used in this study. We upscale the original H&E image by a factor of 1.33 using Lanczos-based resampling ^1^ to approximate single-cell resolution. This definition ensures that each nucleus is fully contained within the patch while avoiding excessive surrounding tissue. We selected the patch size such that a nucleus with a radius of *r* = 18*µ*m remains entirely visible within a crop. Finally, we remove low-quality Xenium cells exhibiting fewer transcripts than the median transcripts per cell minus the median absolute deviation (Heumos et al., 2023). We implement five-fold spatial cross-validation, where the image is split into 5 evenly sized segments. For each fold, one segment is held out as the test set, while the remaining four segments (80%) form the training set. For each evaluation metric, we collect the five scores, i.e. one from each held-out test fold, and then report their mean and standard deviation. Furthermore, within each training set, we randomly reserve 20% of the cells for validation during H&Enium alignment training.

### 3.3. Unaligned Results

We first evaluate performance on the cell type classification task on frozen foundation model embeddings *z*_*I*_ and *z*_*G*_. Table 1 shows that the pathology foundation models, namely UNI2 and CONCH, surpass the morphological and majority voting baselines. UNI2 beats CONCH on all three slides according to the F1 score, with F1 ranging from 0.52 (BreastILC) to 0.74 (BreastIDC). Taking into account UNI2’s superior performance, we selected it as our foundation model for the H&Enium alignment.

**Table 1.**
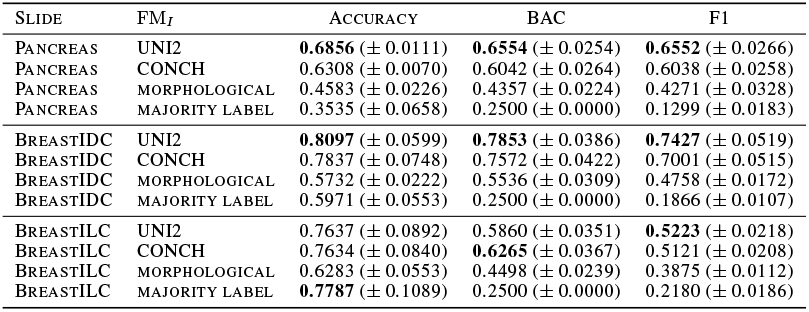
Cell type classification performance across five spatial folds for pathology foundation models (FM_*I*_) versus baselines. Accuracy, balanced accuracy (BAC), and F1 score (F1) are reported as mean *±* standard deviation.

Appendix Table 7 shows that transcriptomic foundation models (CellPLM and scGPT) and the raw gene expression baseline (*G*) achieve high cell type classification scores, with mean F1 exceeding 0.8 across all slides. On the Breast slides, scGPT and CellPLM yield comparable results (difference in F1 *<* 1%), whereas on the Pancreas slide, CellPLM clearly outperforms scGPT (5% F1 improvenment). We therefore adopt CellPLM as our primary transcriptomic foundation model for the H&Enium alignment.

Comparing Tables 1 and 7 we find that the average performance of models trained for cell type prediction on pathology-derived embeddings is, F1: 0.6401 ([0.5223-0.7427]) across all three slides. The average performance of models trained on gene expression embeddings performed substantially better, F1: 0.9041 ([0.8601-0.9414]). This is highlighting the potential of the aligned latent space 𝒜_*I*_ to better capture informative features from both modalities. Figure 2 additionally shows that in the UMAP (McInnes et al., 2018) visualization, CellPLM produces more coherent cell-type clustering than UNI2.

**Figure 2.**
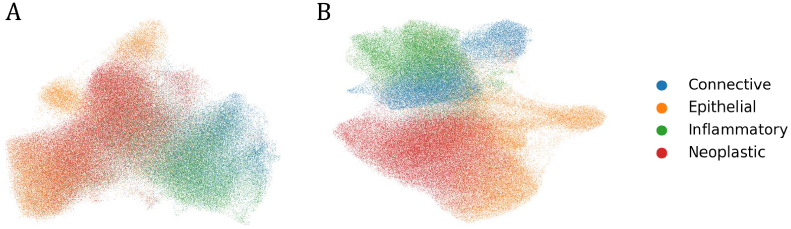
UMAP (McInnes et al., 2018) visualization of Pancreas data colored by cell type, comparing UNI2 (A) and CellPLM (B) single cell embeddings.

**Figure 3.**
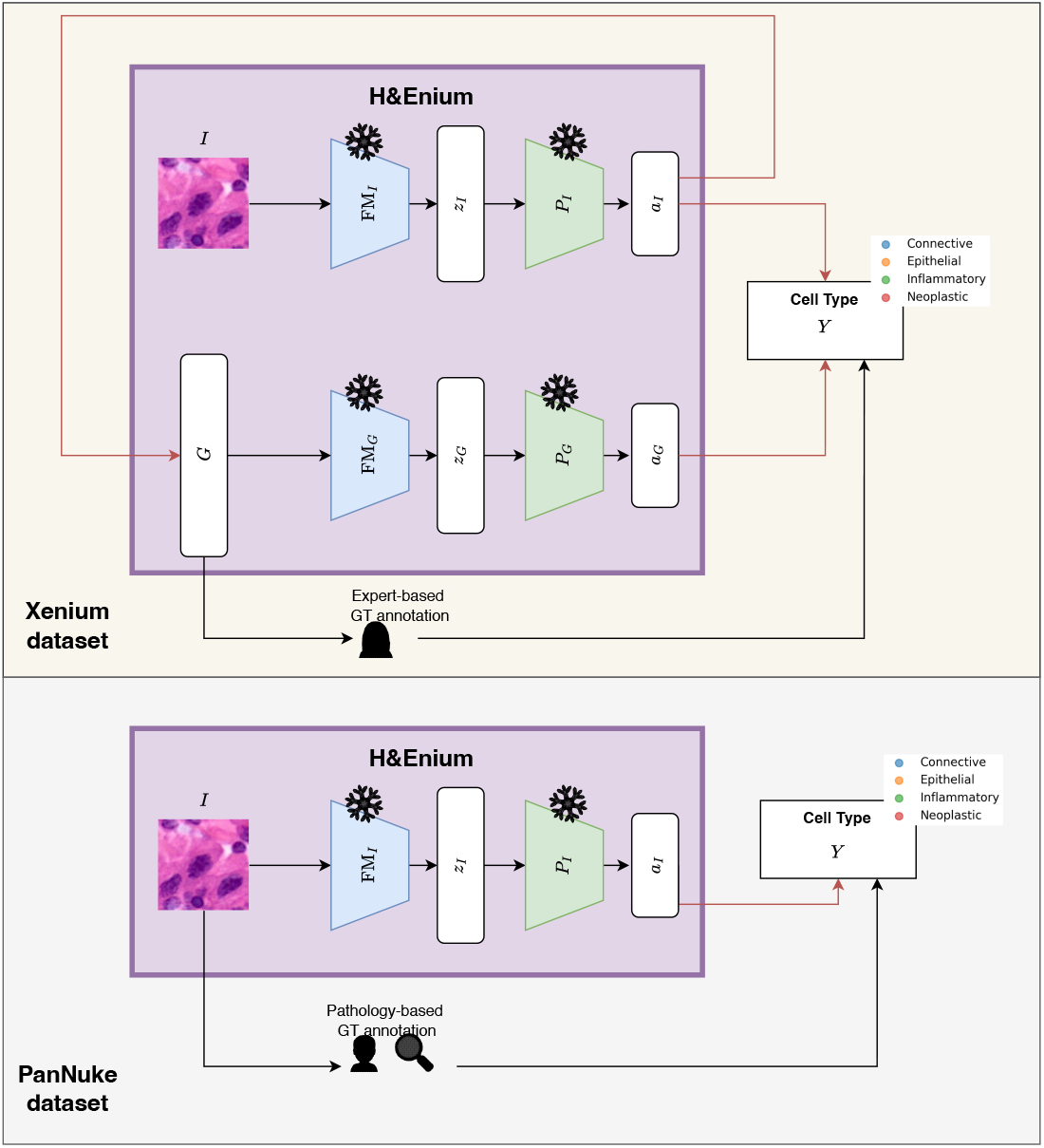
Overview of the H&Enium datasets and their downstream tasks after alignment training is complete (i.e., all H&Enium models are frozen). For the Xenium dataset, we predict the cell type *Y* separately from the image embedding *a*_*I*_ and the gene expression embedding *a*_*G*_. We also predict gene expression *G* from *a*_*I*_. In the PanNuke dataset, only H&E data is available, so we only predict cell type *Y* from *a*_*I*_. Importantly, *Y* is ground truth (GT) annotated directly from the imaging data by an experienced pathologist, whereas in the Xenium dataset, the GT annotations are derived from gene expression *G* and are performed by an expert.

### 3.4. Aligned Results

We choose the best performing foundation models (UNI2 and CellPLM) from Section 3.3 and align their embeddings using H&Enium. Table 2 shows that alignment via our H&Enium model consistently outperforms the non-aligned embeddings when BLEEP_input_ or CLIP are used as targets, whereas using BLEEP as target does not outperform the baseline. The largest relative F1 improvement occurs for BreastILC (*≈* 7%), followed by Pancreas (*≈* 2%), while BreastIDC shows gains under 1%.

**Table 2.**
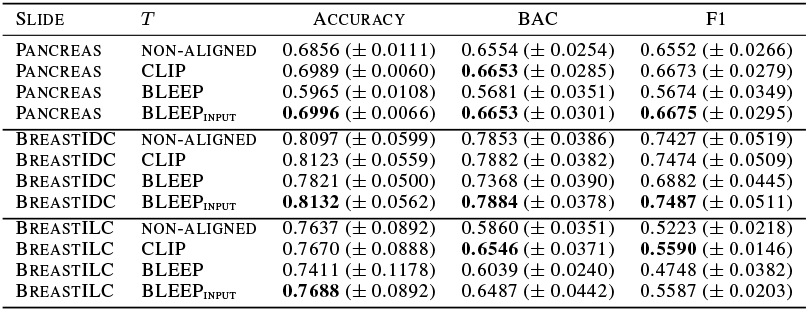
Cell type classification performance across pathology foundation models including non-aligned baseline and H&Enium aligned models. Metrics as mean *±* standard deviation.

| SLIDE | $T$ | ACCURACY | BAC | F1 |
| --- | --- | --- | --- | --- |
| PANCREAS | NON-ALIGNED | 0.6856 (± 0.0111) | 0.6554 (± 0.0254) | 0.6552 (± 0.0266) |
| PANCREAS | CLIP | 0.6989 (± 0.0060) | **0.6653** (± 0.0285) | 0.6673 (± 0.0279) |
| PANCREAS | BLEEP | 0.5965 (± 0.0108) | 0.5681 (± 0.0351) | 0.5674 (± 0.0349) |
| PANCREAS | BLEEP<sub>INPUT</sub> | **0.6996** (± 0.0066) | **0.6653** (± 0.0301) | **0.6675** (± 0.0295) |
| BREASTIDC | NON-ALIGNED | 0.8097 (± 0.0599) | 0.7853 (± 0.0386) | 0.7427 (± 0.0519) |
| BREASTIDC | CLIP | 0.8123 (± 0.0559) | 0.7882 (± 0.0382) | 0.7474 (± 0.0509) |
| BREASTIDC | BLEEP | 0.7821 (± 0.0500) | 0.7368 (± 0.0390) | 0.6882 (± 0.0445) |
| BREASTIDC | BLEEP<sub>INPUT</sub> | **0.8132** (± 0.0562) | **0.7884** (± 0.0378) | **0.7487** (± 0.0511) |
| BREASTILC | NON-ALIGNED | 0.7637 (± 0.0892) | 0.5860 (± 0.0351) | 0.5223 (± 0.0218) |
| BREASTILC | CLIP | 0.7670 (± 0.0888) | **0.6546** (± 0.0371) | **0.5590** (± 0.0146) |
| BREASTILC | BLEEP | 0.7411 (± 0.1178) | 0.6039 (± 0.0240) | 0.4748 (± 0.0382) |
| BREASTILC | BLEEP<sub>INPUT</sub> | **0.7688** (± 0.0892) | 0.6487 (± 0.0442) | 0.5587 (± 0.0203) |

We also assess cell type classification performance of Xenium-trained models on the out-of-sample PanNuke data, whose H&E images are annotated by expert pathologists. Specifically, we apply Pancreas-trained models to the PanNuke Pancreas subset and Breast-trained models to the PanNuke Breast subset. We compare H&Enium aligned models with BLEEP_input_, CLIP as targets to the non-aligned baseline in Table 3. Alignment with BLEEP_input_ delivers more than 16% relative improvement in F1 for Pancreas, BreastIDC and BreastILC.

**Table 3.**
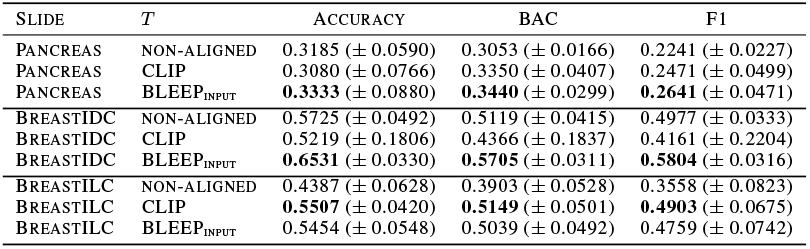
Cell type classification performance on out-of-sample PanNuke dataset across pathology foundation models including nonaligned baseline and H&Enium aligned models. Metrics as mean *±* standard deviation. Evaluation is performed on the entire test set using the five models trained on spatial Xenium folds.

| SLIDE | $T$ | ACCURACY | BAC | F1 |
| --- | --- | --- | --- | --- |
| PANCREAS | NON-ALIGNED | 0.3185 (± 0.0590) | 0.3053 (± 0.0166) | 0.2241 (± 0.0227) |
| PANCREAS | CLIP | 0.3080 (± 0.0766) | 0.3350 (± 0.0407) | 0.2471 (± 0.0499) |
| PANCREAS | BLEEP<sub>INPUT</sub> | **0.3333** (± 0.0880) | **0.3440** (± 0.0299) | **0.2641** (± 0.0471) |
| BREASTIDC | NON-ALIGNED | 0.5725 (± 0.0492) | 0.5119 (± 0.0415) | 0.4977 (± 0.0333) |
| BREASTIDC | CLIP | 0.5219 (± 0.1806) | 0.4366 (± 0.1837) | 0.4161 (± 0.2204) |
| BREASTIDC | BLEEP<sub>INPUT</sub> | **0.6531** (± 0.0330) | **0.5705** (± 0.0311) | **0.5804** (± 0.0316) |
| BREASTILC | NON-ALIGNED | 0.4387 (± 0.0628) | 0.3903 (± 0.0528) | 0.3558 (± 0.0823) |
| BREASTILC | CLIP | **0.5507** (± 0.0420) | **0.5149** (± 0.0501) | **0.4903** (± 0.0675) |
| BREASTILC | BLEEP<sub>INPUT</sub> | 0.5454 (± 0.0548) | 0.5039 (± 0.0492) | 0.4759 (± 0.0742) |

Further, we analyze gene expression prediction of aligned latent space embeddings in Table 4. Aligned embeddings with targets BLEEP_input_ and CLIP consistently outperform the unaligned baseline in PCC and RVD across all slides. For BreastILC, BreastIDC and Pancreas, the relative increases in PCC exceed 10%.

**Table 4.** Gene expression prediction performance across pathology foundation models non-aligned baseline and H&Enium aligned models. Metrics as mean *±* standard deviation.

| SLIDE | $T$ | PCC ↑ | RVD ↓ |
| --- | --- | --- | --- |
| PANCREAS | NON-ALIGNED | 0.3612 (± 0.0335) | 0.7283 (± 0.0174) |
| PANCREAS | CLIP | **0.4014** (± 0.0316) | **0.6603** (± 0.0295) |
| PANCREAS | BLEEP | 0.3297 (± 0.0393) | 0.7589 (± 0.0233) |
| PANCREAS | BLEEP<sub>INPUT</sub> | 0.4012 (± 0.0316) | 0.6617 (± 0.0274) |
| BREASTIDC | NON-ALIGNED | 0.4313 (± 0.0117) | 0.6323 (± 0.0257) |
| BREASTIDC | CLIP | 0.4754 (± 0.0089) | **0.5428** (± 0.0161) |
| BREASTIDC | BLEEP | 0.4306 (± 0.0099) | 0.6258 (± 0.0193) |
| BREASTIDC | BLEEP<sub>INPUT</sub> | **0.4755** (± 0.0085) | 0.5480 (± 0.0127) |
| BREASTILC | NON-ALIGNED | 0.3337 (± 0.0149) | 0.7437 (± 0.0155) |
| BREASTILC | CLIP | 0.3688 (± 0.0116) | **0.6733** (± 0.0303) |
| BREASTILC | BLEEP | 0.3190 (± 0.0124) | 0.7591 (± 0.0231) |
| BREASTILC | BLEEP<sub>INPUT</sub> | **0.3698** (± 0.0113) | 0.6744 (± 0.0300) |

## 4. Discussion and Conclusion

The main contributions of this work are: (1) the adaptation of patch-level pathology foundation models to single cells, (2) the introduction of a novel soft alignment target for cross-modal embedding alignment, (3) a demonstration of improved cell type classification and gene expression prediction on independent H&E datasets.

Our aligned representation yields richer H&E-based embeddings that substantially improve both cell-type classification and gene-expression prediction. In out-of-sample evaluations, models trained on these embeddings showed increased performance of 16% on cell type prediction and 10% on gene expression prediction. To our knowledge this work is the first single-cell level alignment of the H&E image and transcriptomics modalities. Future efforts should involve benchmarking its performance against patch-level models in a pseudo-bulk manner. Additionally, this provides a foundation for scaling the training and architecture across various slides and tumor types. Ultimately, H&Enium establishes a framework for aligning pathology and transcriptomics at single-cell resolution, enhancing the potential of H&E-only analysis pipelines in both research and clinical settings.

## A. Soft Cross Entropy Definition

We define the softmax function with learnable temperature *τ >* 0 row-wise over a matrix *X* ∈ ℝ^*B×B*^ as:

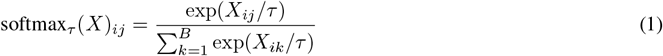

We define a soft cross-entropy function, which operates row-wise over logits and targets:

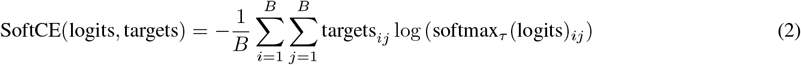

This function is equivalent to applying standard cross-entropy with soft probability targets.^2^

## B. Target definitions

We experiment with different strategies for defining the (soft) target matrix *T*. Each approach encodes different assumptions about inter-sample relationships.

### CLIP Target

The CLIP-style target (Radford et al., 2021) assumes a strict one-to-one correspondence between image and gene embeddings within a batch. It uses an identity matrix as the target distribution:

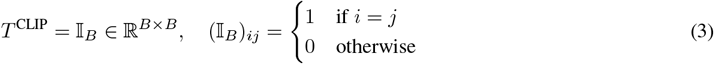

This enforces hard alignment between matched pairs only.

### BLEEP Target

The BLEEP target introduces soft alignment by measuring intra-modality similarity over the output embeddings (Xie et al., 2023). This is advantageous because, within each batch, gene and image embeddings often exhibit significant similarity, particularly among cells of the same type. BLEEP therefore computes a weighted combination of intra-image and intra-gene similarities:

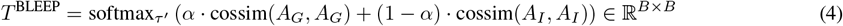

Here, *τ* ^*′*^ is a softmax temperature hyperparameter (non-learnable), and *α* ∈ [0, 1] controls the relative weight of gene vs. image similarity.

### BLEEP_input_ Target

We extend BLEEP by computing intra-modality similarities based on *Z* instead of *A* since we consider the pre-projection embeddings from the foundation models to be more reliable, especially at the early stages of H&Enium model training.

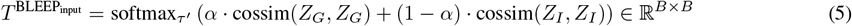

## C. Projection Heads

*P*_*I*_ and *P*_*G*_ are simple one-layer projections with *d*_*a*_ = 128 and batch size *B* = 64 using GELU activation (Hendrycks & Gimpel, 2016), Layer Normalization (Ba et al., 2016) and dropout (Srivastava et al., 2014) with *p* = 0.3 using the AdamW optimizer (Loshchilov & Hutter, 2017) with an initial learning rate of 0.001 and a weight decay of 0.0001, training for a maximum of 20 epochs with early stopping after 5 epochs, only saving model checkpoints if validation loss decreases.

## D. Relative Variation Distance

Relative Variation Distance (RVD) is defined as follows:

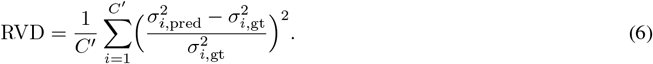

*C*^*′*^ denotes the number of predicted genes (*C*^*′*^ = 50 for this work) and 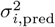 is the variance of the predicted expression for gene *i* across cells, and 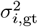 is the variance of the true expression for gene *i* across cells.

The RVD metric was introduced in Zhu et al. (2025) in response to Xie et al. (2023)’s observation that in log-transformed space, a naive baseline predicting the cell-wise mean expression across all genes can produce deceptively high PCC. RVD thus quantifies the average squared relative deviation between predicted and true gene variances, providing a more sensitive measure of how well the model captures heterogeneity in gene expression across cells.

## E. Morphological Baseline

Xenium provides cell segmentation information that allows us to derive features based on morphology. In contrast to H&E images - which capture detailed visual cues such as color, texture, and tissue architecture - the features presented here are solely derived from the geometric shapes of cells and nuclei and their neighbors. This morphological baseline serves as a comparison to the features extracted from pathology foundation models.

Every cell is represented by a set of vertices that define its boundary, and each cell is associated with a nucleus that is also defined by its own vertices. These vertex coordinates provide the necessary information to calculate geometric descriptors that characterize the shape and spatial relationships of the cells. Based on the segmentation information, we compute 16 features:

- **Cell Features:**

- **Cell Area:** The area enclosed by the cell boundary.
- **Cell Maximum Radius:** The maximum distance from the cell centroid to the cell boundary.
- **Cell Perimeter:** The total length of the cell boundary.
- **Cell Perimeter-to-Area Ratio:** A measure that reflects the compactness of the cell shape.
- **Cell Concavity:** An indicator of the deviation of the cell shape from a perfect circle.
- **Cell Smoothness:** Quantifying the regularity of the cell boundary, i.e. the perimeter divided by the number of boundary vertices.

- **Nucleus Features:**

- **Nucleus Area:** The area enclosed by the nucleus boundary.
- **Nucleus Maximum Radius:** The maximum distance from the nucleus centroid to its boundary.
- **Nucleus Perimeter:** The total length of the nucleus boundary.
- **Nucleus Perimeter-to-Area Ratio:** A descriptor of the nucleus shape.
- **Nucleus Concavity:** A measure of the deviation of the nucleus shape from an ideal circle.

- **Combined Features:**

- **Nucleus-to-Cell Area Ratio:** The ratio of the nucleus area to the cell area.
- **Nucleus-to-Cell Centroid Distance:** The Euclidean distance between the centroids of the cell and its nucleus.
- **Nearest-Nucleus Features:** Spatial features based on a nearest neighbor search:

* The distance from the nucleus centroid to the nearest nucleus centroid.
* The distance from the cell membrane to the nearest nucleus centroid.
* The distance from the cell membrane to the *k*th nearest nucleus centroid.

## F. Datasets overview

### F.1. Pancreas

The Xenium Pancreas dataset comprises 190,965 cells and employs a gene panel of size *C* = 474. After preprocessing, 147,707 cells are left for training and testing, see Table 5 for cell type distribution statistics across folds. Downloaded from the 10x Genomics Xenium dataset page.

**Table 5.** Distribution of cell types across training and testing folds for each Xenium slide.

| SLIDE | FOLD | SET | NUM. CELLS | CELL TYPES $Y$ | | | |
| --- | --- | --- | --- | --- | --- | --- | --- |
|  |  |  |  | CONNECTIVE | INFLAMMATORY | NEOPLASTIC | EPITHELIAL |
| PANCREAS | OVERALL |  | 147707 | 30470 (20.63%) | 29253 (19.80%) | 52208 (35.35%) | 35776 (24.22%) |
| PANCREAS | 0 | TRAIN | 118166 | 23476 (19.87%) | 22375 (18.94%) | 44689 (37.82%) | 27626 (23.38%) |
| PANCREAS | 0 | TEST | 29541 | 6994 (23.68%) | 6878 (23.28%) | 7519 (25.45%) | 8150 (27.59%) |
| PANCREAS | 1 | TRAIN | 118166 | 23606 (19.98%) | 23417 (19.82%) | 41768 (35.35%) | 29375 (24.86%) |
| PANCREAS | 1 | TEST | 29541 | 6864 (23.24%) | 5836 (19.76%) | 10440 (35.34%) | 6401 (21.67%) |
| PANCREAS | 2 | TRAIN | 118166 | 27131 (22.96%) | 25615 (21.68%) | 39288 (33.25%) | 26132 (22.11%) |
| PANCREAS | 2 | TEST | 29541 | 3339 (11.30%) | 3638 (12.32%) | 12920 (43.74%) | 9644 (32.65%) |
| PANCREAS | 3 | TRAIN | 118166 | 25615 (21.68%) | 24483 (20.72%) | 41149 (34.82%) | 26919 (22.78%) |
| PANCREAS | 3 | TEST | 29541 | 4855 (16.43%) | 4770 (16.15%) | 11059 (37.44%) | 8857 (29.98%) |
| PANCREAS | 4 | TRAIN | 118164 | 22052 (18.66%) | 21122 (17.88%) | 41938 (35.49%) | 33052 (27.97%) |
| PANCREAS | 4 | TEST | 29543 | 8418 (28.49%) | 8131 (27.52%) | 10270 (34.76%) | 2724 (9.22%) |
| BREASTIDC | OVERALL |  | 439534 | 89125 (20.28%) | 66850 (15.21%) | 262434 (59.71%) | 21125 (4.81%) |
| BREASTIDC | 0 | TRAIN | 351628 | 73702 (20.96%) | 55667 (15.83%) | 202771 (57.67%) | 19488 (5.54%) |
| BREASTIDC | 0 | TEST | 87906 | 15423 (17.54%) | 11183 (12.72%) | 59663 (67.87%) | 1637 (1.86%) |
| BREASTIDC | 1 | TRAIN | 351628 | 69414 (19.74%) | 54081 (15.38%) | 210515 (59.87%) | 17618 (5.01%) |
| BREASTIDC | 1 | TEST | 87906 | 19711 (22.42%) | 12769 (14.53%) | 51919 (59.06%) | 3507 (3.99%) |
| BREASTIDC | 2 | TRAIN | 351628 | 71433 (20.31%) | 54163 (15.40%) | 207859 (59.11%) | 18173 (5.17%) |
| BREASTIDC | 2 | TEST | 87906 | 17692 (20.13%) | 12687 (14.43%) | 54575 (62.08%) | 2952 (3.36%) |
| BREASTIDC | 3 | TRAIN | 351628 | 70215 (19.97%) | 48622 (13.83%) | 213845 (60.82%) | 18946 (5.39%) |
| BREASTIDC | 3 | TEST | 87906 | 18910 (21.51%) | 18228 (20.74%) | 48589 (55.27%) | 2179 (2.48%) |
| BREASTIDC | 4 | TRAIN | 351624 | 71736 (20.40%) | 54867 (15.60%) | 214746 (61.07%) | 10275 (2.92%) |
| BREASTIDC | 4 | TEST | 87910 | 17389 (19.78%) | 11983 (13.63%) | 47688 (54.25%) | 10850 (12.34%) |
| BREASTILC | OVERALL |  | 270700 | 34192 (12.63%) | 23437 (8.66%) | 210800 (77.87%) | 2271 (0.84%) |
| BREASTILC | 0 | TRAIN | 216560 | 20712 (9.56%) | 16185 (7.47%) | 178820 (82.57%) | 843 (0.39%) |
| BREASTILC | 0 | TEST | 54140 | 13480 (24.90%) | 7252 (13.39%) | 31980 (59.07%) | 1428 (2.64%) |
| BREASTILC | 1 | TRAIN | 216560 | 27677 (12.78%) | 18565 (8.57%) | 168204 (77.67%) | 2114 (0.98%) |
| BREASTILC | 1 | TEST | 54140 | 6515 (12.03%) | 4872 (9.00%) | 42596 (78.68%) | 157 (0.29%) |
| BREASTILC | 2 | TRAIN | 216560 | 29882 (13.80%) | 20517 (9.47%) | 164034 (75.75%) | 2127 (0.98%) |
| BREASTILC | 2 | TEST | 54140 | 4310 (7.96%) | 2920 (5.39%) | 46766 (86.38%) | 144 (0.27%) |
| BREASTILC | 3 | TRAIN | 216560 | 29804 (13.76%) | 19256 (8.89%) | 165411 (76.38%) | 2089 (0.96%) |
| BREASTILC | 3 | TEST | 54140 | 4388 (8.10%) | 4181 (7.72%) | 45389 (83.84%) | 182 (0.34%) |
| BREASTILC | 4 | TRAIN | 216560 | 28693 (13.25%) | 19225 (8.88%) | 166731 (76.99%) | 1911 (0.88%) |
| BREASTILC | 4 | TEST | 54140 | 5499 (10.16%) | 4212 (7.78%) | 44069 (81.40%) | 360 (0.66%) |

### F.2. Breast

We use two publicly available Breast slides from Xenium. The BreastIDC and BreastILC datasets both profile *C* = 380, containing 574,527 and 365,604 cells, respectively. Statistics for cells passing the preprocessing are shown in Table 5. Downloaded from the 10x Genomics Xenium dataset page.

### F.3. PanNuke

Out-of-sample test data from the PanNuke dataset (Gamper et al., 2020) is downloaded from the Warwick Tissue Image Analytics (TIA) Centre. For Pancreas, 741 single-cell image patches are available, Breast contains 8471 cells. Table 6 shows the respective cell type distributions.

**Table 6.** PanNuke cell type distribution on the PanNuke data used for out-of-sample testing.

| ORGAN | NUM. CELLS | CELL TYPES $Y$ | | | |
| --- | --- | --- | --- | --- | --- |
|  |  | CONNECTIVE | INFLAMMATORY | NEOPLASTIC | EPITHELIAL |
| PANCREAS | 741 | 394 (53.17%) | 146 (19.70%) | 127 (17.14%) | 74 (9.99%) |
| BREAST | 8471 | 1826 (21.56%) | 1057 (12.48%) | 3191 (37.67%) | 2397 (28.30%) |

### F.4. Architecture and Datasets

## G. Cell Type Classification for Transcriptomic Foundation Models

**Table 7.** Cell type classification performance across five spatial folds for transcriptomic foundation models (FM_*G*_) versus baselines. Accuracy, balanced accuracy (BAC), and F1 score (F1) are reported as mean *±* standard deviation.

| SLIDE | $FM_G$ | ACCURACY | BAC | F1 |
| --- | --- | --- | --- | --- |
| PANCREAS | CELLPLM | 0.8709 (± 0.0071) | 0.8593 (± 0.0065) | 0.8601 (± 0.0046) |
| PANCREAS | SCGPT | 0.8242 (± 0.0178) | 0.8132 (± 0.0136) | 0.8136 (± 0.0118) |
| PANCREAS | GENE EXPRESSION | **0.9170** (± 0.0062) | **0.9081** (± 0.0075) | **0.9080** (± 0.0089) |
| PANCREAS | MAJORITY LABEL | 0.3535 (± 0.0658) | 0.2500 (± 0.0000) | 0.1299 (± 0.0183) |
| BREASTIDC | CELLPLM | 0.9615 (± 0.0226) | 0.9607 (± 0.0130) | 0.9414 (± 0.0225) |
| BREASTIDC | SCGPT | 0.9623 (± 0.0211) | **0.9610** (± 0.0118) | **0.9442** (± 0.0217) |
| BREASTIDC | GENE EXPRESSION | **0.9631** (± 0.0183) | 0.9603 (± 0.0119) | 0.9439 (± 0.0206) |
| BREASTIDC | MAJORITY LABEL | 0.5971 (± 0.0553) | 0.2500 (± 0.0000) | 0.1866 (± 0.0107) |
| BREASTILC | CELLPLM | 0.9719 (± 0.0133) | **0.9600** (± 0.0097) | 0.9107 (± 0.0059) |
| BREASTILC | SCGPT | **0.9722** (± 0.0120) | 0.9509 (± 0.0093) | **0.9112** (± 0.0115) |
| BREASTILC | GENE EXPRESSION | 0.9720 (± 0.0120) | 0.9456 (± 0.0096) | 0.9046 (± 0.0117) |
| BREASTILC | MAJORITY LABEL | 0.7787 (± 0.1089) | 0.2500 (± 0.0000) | 0.2180 (± 0.0186) |

## Footnotes

1 See the documentation of Pillow resampling filters.

2 Implemented with PyTorch’s cross entropy function

